# Does oxidative stress shorten telomeres *in vivo*? A meta-analysis

**DOI:** 10.1101/2022.08.17.504247

**Authors:** Emma Armstrong, Jelle Boonekamp

## Abstract

Telomere attrition is considered a hallmark of ageing. Untangling the proximate causes of telomere attrition may therefore reveal important aspects about the ageing process. In a landmark paper in 2002 Thomas von Zglinicki demonstrated that oxidative stress causes telomere attrition in cell culture. In the next 20 years, oxidative stress became firmly embedded into modern theories of ageing and telomere attrition. However, a recent surge of *in vivo* studies reveals an inconsistent pattern questioning the unequivocal role of oxidative stress in telomere dynamics, in living organisms. Here we report the results of the first formal meta-analysis on the association between oxidative stress and telomere dynamics *in vivo*, representing 37 studies, 4,834 individuals, and 18,590 correlational measurements. The overall correlation between oxidative stress markers and telomere dynamics was indistinguishable from zero. This result was independent of the type of oxidative stress marker, telomere dynamic, or taxonomic group. However, telomere measurement method affected the analysis with TRF but not qPCR-based studies showing a significant overall correlation. The correlation was more pronounced in short-lived species and during the adult life phase, when ageing becomes apparent. We then performed an additional meta-analysis of interventional studies (n=7) manipulating oxidative stress. This revealed a significant effect of treatment on telomere dynamics. Our findings indicate that oxidative stress may have a profound effect on telomere dynamics in living organisms fundamentally underpinning the process of ageing.

## Introduction

Telomeres comprise noncoding tandem DNA repeats and associated sheltering proteins that form protective envelopes at the chromosome-ends. They are essential for maintaining chromosomal integrity and function (Blackburn 1991). Telomeres progressively shorten with age in somatic tissues, eventually reaching a critical length, triggering cell cycle arrest (Blackburn 1991; Saretzki et al. 1999). Progressive telomere loss through the life course is considered a hallmark of ageing (López-Otín et al. 2013) and appears associated with ageing-related disease (Armanios & Blackburn 2012; Rossiello et al. 2022), life stress (Epel et al. 2004) and mortality risk in humans (Boonekamp et al. 2013). Telomeres are now also increasingly studied in non-model species and in natural populations. These studies show that telomere length is related to sexual ornamentation (Kauzálová et al. n.d.), immune function (Roast et al. 2022), environmental stressors (Chatelain et al. 2019; Eastwood et al. 2022), and mortality risk across a wide range of vertebrate species (Wilbourn et al. 2018). Furthermore, telomere length may reflect phenotypic quality as indicated by the relationship between early life telomere length and longevity seen in captive and wild bird populations (Sheldon et al. 2021; Eastwood et al. 2019; Heidinger et al. 2012). All of this indicates that telomere- and life-history dynamics are entwined, with differing patterns depending on species, the environment, and life stage (Monaghan 2010).

It is of interest to identify the underlying causes of telomere attrition as this may provide crucial insight into the processes of ageing, their connection with the environment, and variation between species with differing life-history strategies. Oxidative stress is thought to be one of the main drivers of telomere attrition (Reichert & Stier 2017; López-Otín et al. 2013; Metcalfe & Olsson 2021). The popularity of this hypothesis is evident in the studies of our meta-analysis. Oxidative stress is the result of the inevitable production of reactive oxygen species (ROS) during energy synthesis by the mitochondria. Oxidative damage occurs when ROS are not sufficiently neutralized by circulating antioxidants leading to oxidative stress (Costantini & Verhulst 2009). Telomeres form a preferential target for ROS as they contain many guanine base pairs prone to oxidation and they are particularly vulnerable when in a single-stranded form during mitosis (Zglinicki et al. 2000). However, the current evidence for the oxidative stress hypothesis for telomere attrition largely hinges on experimental studies demonstrating a causal relationship in cell culture (Zglinicki 2002). While these experiments convincingly show that elevation of oxidative stress accelerates telomere loss *in vitro*, it is unclear whether this means a functional relationship should also exist under natural oxidative stress levels *in vivo* (Boonekamp et al. 2017). There are numerous reasons for questioning this. For example, as previously recognized, it is unclear how to scale the elevated oxidative stress levels that were applied to cells *in vitro* to the naturally occurring levels of oxidative stress in living organisms (Pineda-Pampliega et al. 2020). Furthermore, cells will behave differently when they are part of a physiological system; Organisms may regulate levels of oxidative stress by adjusting their behavior, or they may sacrifice certain tissues to maintain critical functions. The rate of telomere attrition also depends on other factors such as cell proliferation rate. Other causes of telomere shortening include the end-replication problem (the inability of DNA polymerase to replicate the final few bases (Olovnikov 1973)) and erosion of the single strand overhand (Stewart et al. 2003). Telomere shortening may also be under the influence of metabolic processes that are unrelated to oxidative stress (Casagrande & Hau 2019). This means that while oxidative stress may be important, it may not necessarily be the primary cause of telomere attrition *in vivo*. Recently, a growing number of studies measured the relationship between oxidative stress and telomere length or rate of attrition *in vivo*. While some of these studies support this relationship (Pineda-Pampliega et al. 2020), others show no relationship (Boonekamp et al. 2017) or even report an association in the opposite direction (Criscuolo et al. 2019). These inconsistencies challenge the idea of an unequivocal causal relationship between oxidative stress and telomere attrition in living organisms.

Here we conduct the first formal meta-analysis reviewing all the evidence currently available on the correlation between markers of oxidative stress and telomere dynamics for humans and other vertebrate species. We first test if there is a significant overall relationship between markers of oxidative stress and telomere dynamics. We then divide the analysis into sub-sections per type of oxidative stress marker (damage vs. protection) and telomere dynamic (length vs. rate of attrition) to reveal any contrasts. We perform a moderator analysis to investigate the effects of telomere length measurement method, taxonomic group, life-phase, and species longevity. Finally, we report the results of a separate meta-analysis of experimental studies that elevated oxidative damage or antioxidant protection and measured the effects on telomere dynamics.

## Results

### Associative studies

Our meta-analysis included 107 Pearson correlation effect sizes of 37 papers that collectively comprised 18,590 measurements of the relationship between oxidative stress markers and telomere dynamics in 4,834 individuals (Fig. 1). Table 1 provides an overview of the studies that were included in our meta-analysis. The overall Pearson correlation between oxidative stress markers and telomere dynamics was low (*r* = 0.025) and indistinguishable from zero (95% CI - 0.053; 0.103; *p* = 0.53; Fig. 1). Subset-analysis indicated this result is consistent for the associations between telomere length and antioxidants (*r*= 0.031; CI −0.075; 0.138), telomere length and oxidants (*r* = 0.003; CI −0.102; 0.109), telomere shortening and antioxidants (*r* = 0.062; CI −0.011; 0.134), and telomere shortening and oxidants (*r* = 0.001; CI −0.177; 0.179; Fig. 1). Based on these initial findings it appears that oxidative stress and telomere dynamics are unrelated.

**Figure 1.**
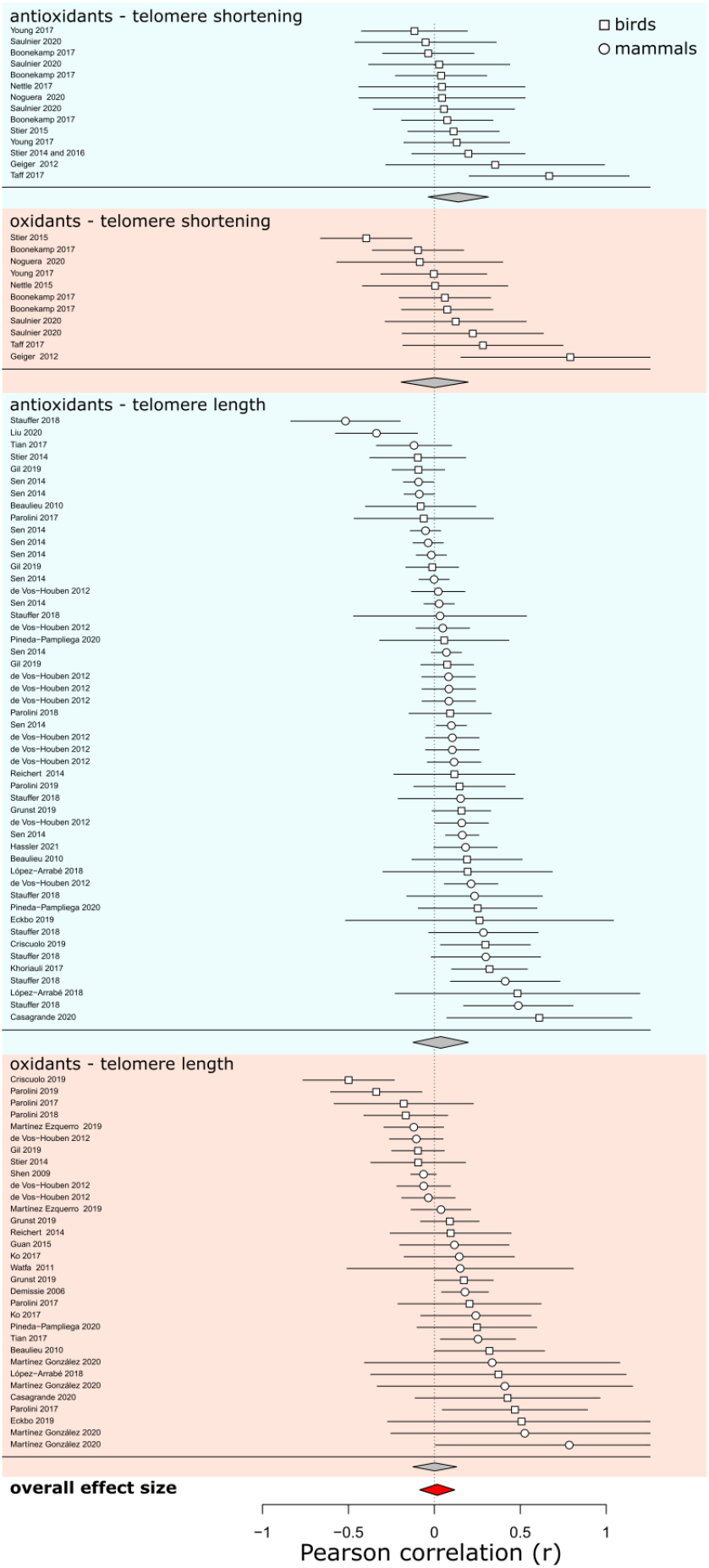
Forest plot showing the Pearson correlations between oxidative stress markers and telomere dynamics. Positive correlations support the hypothesis that oxidative stress causes telomere attrition (see methods for further details on the directionality of effect sizes). Squares and circles indicate the relationships for birds and mammals, respectively. Horizontal bars reflect the 95% confidence intervals. Grey diamonds reflect the summary effect sizes per subset analysis (indicated by the subheadings). The single red diamond at the bottom shows the total overall effect size. Diamond widths reflect their 95% confidence intervals.

**Table 1.** Overview of studies that were included in the associative and experimental treatment effects meta-analyses with the moderator variables class, life phase, longevity, telomere method, and oxidative stress type. Further information shown here is the study species and the specific oxidative stress markers that were used. The effect sizes are available in our online supporting data table information (S1) with corresponding paper ID’s.

| Paper* | Reference | Study type | Treatment | Class | Species | Life phase | Longevity | Telomere dynamic | Telomere method | Oxidative stress type | Oxidative stress marker(s) |
| --- | --- | --- | --- | --- | --- | --- | --- | --- | --- | --- | --- |
| 1 | (Eckbo et al. 2019) | Associative | na | Aves | *Cephus grylle* | Adult | 29.9 | Length | qPCR | Both | TAC, ROM |
| 2 | (Martínez-González et al. 2020) | Associative | na | Mammalia | *Mus musculus* | Young | 4 | Length | qPCR | Damage | ROM, MDA |
| 3 | (Pineda-Pampliega et al. 2020) | Associative | na | Aves | *Ciconia ciconia* | Young | 39 | Length | TRF | Both | MDA, Tocopherol, TAC |
| 4 | (Demissie et al. 2006) | Associative | na | Mammalia | *Homo sapiens* | Adult | 122.5 | Length | TRF | Damage | Urinary 8-epi-PGF2α |
| 5 | (Taff & Freeman-Gallant 2017) | Associative | na | Aves | *Geothlypis trichas* | Adult | 11.5 | Loss | qPCR | Both | TAC, Comet assay |
| 6 | (Geiger et al. 2012) | Associative | na | Aves | *Aptenodytes patagonicus* | Young | 27 | Loss | qPCR | Both | TAC, ROM |
| 7 | (Nettle et al. 2017) | Associative | na | Aves | *Sturnus vulgaris* | Young | 22.9 | Loss | qPCR | Antioxidant | TAC |
| 8 | (Boonekamp et al. 2017) | Associative | na | Aves | *Corvus monedula* | Adult | 20.3 | Loss | TRF | Both | Uric acid, Redox state, GSH, GSSG, dROM, TBARS |
| 9 | (Parolini et al. 2017) | Associative | na | Aves | *Larus michahellis* | Young | 19.2 | Length | qPCR | Both | TOS, TAC, Protein carbonyls, TBARS |
| 10 | (Grunst et al. 2019) | Associative | na | Aves | *Zonotrichia albicollis* | Young | 14.9 | Length | qPCR | Both | TAC, ROM |
| 11 | (Parolini et al. 2019) | Associative | na | Aves | *Larus michahellis* | Young | 19.2 | Length | qPCR | Both | TAC, TOS |
| 12 | (Stauffer et al. 2018) | Associative | na | Mammalia | *Mus musculus* | Young | 4 | Length | qPCR | Antioxidant | SOD, CAT, GP, GR, G6PDH, GSH/GSSG, GST |
| 13 | (López-Arrabé et al. 2018) | Associative | na | Aves | *Ficedula hypoleuca* | Adult | 15 | Length | qPCR | Both | MDA, TAS, GSH |
| 14 | (Parolini et al. 2018) | Associative | na | Aves | *Larus michahellis* | Young | 19.2 | Length | qPCR | Both | TAC, TOS |
| 15 | (Khoraiuli et al. 2017) | Associative | na | Aves | *Hirundo rustica* | Adult | 16 | Length | qPCR | Antioxidant | TAC |
| 16 | (Vos-Houben et al. 2012) | Associative | na | Mammalia | *Homo sapiens* | Adult | 122.5 | Length | qPCR | Both | ROM, MDA, GGT, Vitamins, endogenous antioxidants |
| 17 | (Ko et al. 2017) | Associative | na | Mammalia | *Homo sapiens* | Adult | 122.5 | Length | TRF | Damage | MDA, Comet assay |
| 18 | (Guan et al. 2015) | Associative | na | Mammalia | *Homo sapiens* | Adult | 122.5 | Length | TRF | Damage | Urine 8-iso-PGF2α |
| 19 | (Liu et al. 2020) | Associative | na | Mammalia | *Homo sapiens* | Adult | 122.5 | Length | qPCR | Antioxidant | SOD |
| 20 | (Martínez-Ezquerro et al. 2020) | Associative | na | Mammalia | *Homo sapiens* | Adult | 122.5 | Length | qPCR | Damage | DCFH, MDA |
| 21 | (Tian et al. 2017) | Associative | na | Mammalia | *Homo sapiens* | Adult | 122.5 | Length | qPCR | Both | ROM, TAC |
| 22 | (Wafar et al. 2011) | Associative | na | Mammalia | *Homo sapiens* | Adult | 122.5 | Length | TRF | Damage | Carbonyl proteins |
| 23 | (Reichert et al. 2014) | Associative | na | Aves | *Taeniopygia guttata* | Adult | 12 | Length | qPCR | Both | ROM, TAC |
| 24 | (Stier et al. 2015) | Associative | na | Aves | *Parus major* | Young | 15.4 | Loss | qPCR | Both | ROM, TAC |
| 25 | (Stier, Vblanc, et al. 2014) | Associative | na | Aves | *Aptenodytes patagonicus* | Young | 27 | Length | qPCR | Both | ROM, TAC |
| 26 | (Shen et al. 2009) | Associative | na | Mammalia | *Homo sapiens* | Adult | 122.5 | Length | qPCR | Damage | 15-F2t-Isoprostane |
| 27 | (Criscuolo et al. 2019) | Associative | na | Aves | *Sturnus vulgaris* | Young | 22.9 | Length | qPCR | Both | ROM, OXY |
| 28 | (Young et al. 2017) | Associative | na | Aves | *Rissa tridactyla* | Young | 28.5 | Loss | TRF | Both | TAC, Uric acid, Protein carbonyl |
| 29 | (Nettle et al. 2015) | Associative | na | Aves | *Sturnus vulgaris* | Young | 22.9 | Loss | qPCR | Damage | MDA |
| 30 | (Casagrande et al. 2020) | Associative | na | Aves | *Parus major* | Young | 15.4 | Length | TRF | Both | OXY, ROM |
| 31 | (Gil et al. 2019) | Associative | na | Aves | *Sturnus unicolor* | Young | 22.9 | Length | qPCR | Both | GSH, MDA, Uric acid, TAC |
| 32 | (Noguera & Velando 2020) | Associative | na | Aves | *Larus michahellis* | Young | 19.2 | Loss | qPCR | Both | TAC, TBARS |
| 33 | (Saulnier et al. 2020) | Associative | na | Aves | *Taeniopygia guttata* | Adult | 12 | Loss | qPCR | Both | GPx, ROM, GSH, GSSG, unbound GSH |
| 34 | (Sen et al. 2014) | Associative | na | Mammalia | *Homo sapiens* | Adult | 122.5 | Length | qPCR | Antioxidant | Vitamins |
| 35 | (Stier et al. 2014 & 2016) | Associative | na | Aves | *Periparus ater* | Young | 9.5 | Loss | qPCR | Antioxidant | OXY |
| 36 | (Hassler et al. 2021) | Associative | na | Mammalia | *Homo sapiens* | Adult | 122.5 | Length | qPCR | Antioxidant | TAC |
| 37 | (Beaulieu et al. 2010) | Associative | na | Aves | *Pygoscelis adeliae* | Adult | 20 | Length | qPCR | Both | ROM, OXY, Uric acid |
| 3 | (Pineda-Pampliega et al. 2020) | Treatment | Tocopherol + selenium | Aves | *Ciconia ciconia* | Young | 22 | Length | TRF | Na | Na |
| 38 | (Cattan et al. 2008) | Treatment | BSO | Mammalia | *Mus musculus* | Young | 6 | Length | TRF | Na | Na |
| 39 | (Badás et al. 2015) | Treatment | Tocopherol | Aves | *Cyanistes caeruleus* | Young | 3 | Loss | qPCR | Na | Na |
| 40 | (Noguera et al. 2015) | Treatment | Vitamins | Aves | *Taeniopygia guttata* | Adult | 5 | Loss | qPCR | Na | Na |
| 9 | (Parolini et al. 2017) | Treatment | Vitamin E | Aves | *Larus michahellis* | Adult | 12 | Length | qPCR | Na | Na |
| 41 | (Pérez-Rodríguez et al. 2019) | Treatment | Vitamin E, Diquat | Aves | *Ficedula hypoleuca* | Young | 15 | Loss | qPCR | Na | Na |
| 42 | (Kim & Velando 2015) | Treatment | Vitamins | Aves | *Larus michahellis* | Young | 19.2 | Length | qPCR | Na | Na |

\* Effect size calculation was based on the data combined from these two studies.

We found evidence for publication bias (Kendall’s tau = 0.19; p = 0.004). We have previously observed a stronger publication bias for studies that measured telomeres with quantitative Polymerase Chain Reaction (qPCR) in a meta-analysis on the relationship between telomere length and mortality (Wilbourn et al. 2018). We therefore tested for publication bias in the subset of qPCR studies and compared this with the subset of studies that measured telomeres with Terminal Restriction Fragment analysis (TRF), but similar publication biases were present in both subsets (qPCR: Kendall’s tau = 0.19; *p* = 0.01, vs. TRF: Kendall’s tau = 0.34; *p* = 0.02). Publication bias can be problematic because it may inflate the meta-analysis estimate of the true population effect size, but since our overall effect size was close to zero, any bias that may have been present was likely weak.

There was significant heterogeneity among the study effect sizes (*Q* = 225, *p* < .0001) which prompted us to investigate whether there were any meaningful moderator variables underpinning this variation. To this end we tested the effects of the study telomere dynamic (length versus longitudinal rate of loss), the type of oxidative stress marker (damage versus antioxidant), the telomere length measurement method that was used (TRF versus qPCR), taxonomic group (mammal versus bird), the life phase in which the measurements took place (young versus adult), and the natural logarithm of longevity of the study species. Oxidative stress marker type and taxonomic group did not significantly affect the analysis (Fig. 2; Table 2). Somewhat unexpectedly, studies that measured telomere length showed a marginally significantly stronger correlation compared with studies that measured longitudinal telomere loss (Fig. 2; Table 2). Telomere measurement method, life phase, and longevity significantly affected the meta-analysis (all *p*-values < 0.001); The correlation between oxidative stress and telomere dynamics was weaker for studies using young versus adult subjects and for studies using species with greater longevity (Fig.2; Table 2). TRF-based studies showed a stronger correlation between oxidative stress and telomere dynamics compared with studies that measured telomere dynamics with qPCR (Fig. 2; Table 2). The residual heterogeneity remained significant but was substantially reduced by the inclusion of these moderators (*Q* = 179, *p* < 0.001).

**Figure 2.**
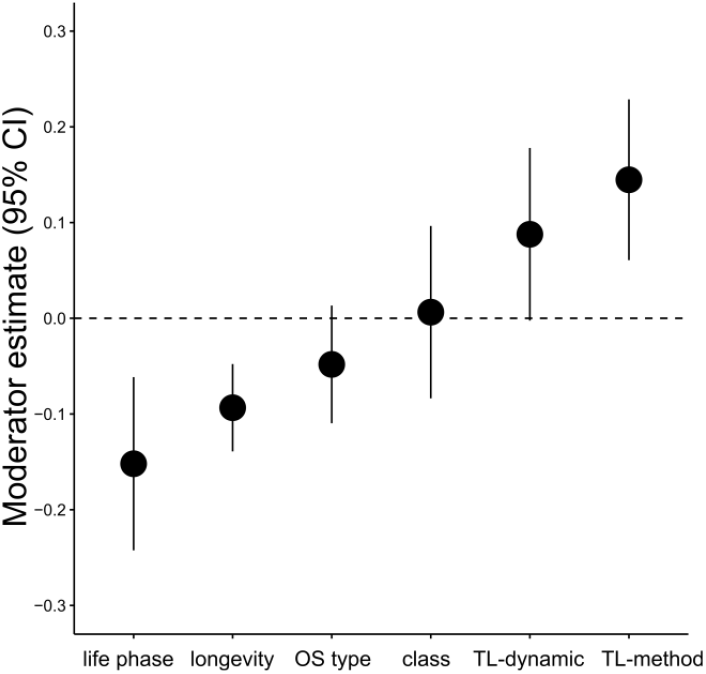
Overview of moderator estimates in the meta-analysis of correlational studies. Error bars represent 95% confidence intervals and deviation from the dotted line therefore represents a significant effect of the moderator. The correlation between oxidative stress and telomere dynamics was significantly weaker both for studies using young versus adult subjects and studies using species with greater longevity. The telomere length dynamic moderator showed a marginally significant trend. Studies that measured telomere length with the TRF method showed a significantly stronger correlation than studies that measured telomere length with qPCR.

**Table 2.** Overview of moderator variables in the meta-analysis of associative studies. The estimates reflect the mean differences between the moderator contrasts as indicated in the moderator description. *P*-values were obtained from z-tests. See methods for further details.

| Moderator | Estimate (95 % confidence interval) | *P*-values |
| --- | --- | --- |
| Telomere dynamic (length vs. attrition) | 0.088 (-0.002; 0.178) | 0.056 |
| Oxidative stress marker type (oxidant vs. antioxidant) | -0.048 (-0.110; 0.013) | 0.125 |
| Telomere measurement method (TRF vs. qPCR) | 0.145 (0.061; 0.229) | <0.001 |
| Life phase (young vs. adult) | -0.152 (-0.242; -0.062) | 0.001 |
| Taxonomic group (Mammalia vs. Aves) | 0.006 (-0.084; 0.096) | 0.889 |
| Species longevity (natural logarithm) | -0.093 (-0.139; -0.048) | <0.001 |

An interesting result of the moderator analysis is that the TRF-based studies showed a significantly stronger association between oxidative stress and telomere dynamics. The TRF method has a higher measurement repeatability (Kärkkäinen et al. 2021) and therefore these studies could potentially reflect a more accurate estimation of the association between oxidative stress and telomere dynamics. We therefore ran a separate meta-analysis for the subset of TRF studies, which confirmed a significant positive correlation between oxidative stress and telomere dynamics (*r* = 0.086; CI 0.012; 0.159; *p* = 0.023). The subset of qPCR studies on the other hand showed a weak overall correlation that was indistinguishable from zero (*r* = 0.02; CI −0.069; 0.109; *p* = 0.662).

### Experimental studies

We performed a separate meta-analysis on 7 experimental studies that measured the effect of oxidative stress interventions on telomere dynamics *in vivo*. See table 1 for an overview of the experimental studies that were included. There was a significant overall effect of treatment on telomere dynamics supporting the hypothesis that oxidative stress causes telomere attrition (Cohen’s *d* = 0.167, 95% CI 0.089; 0.246; *p* < 0.001; Fig. 3). There was no evidence for publication bias (Kendall’s tau = 0.52, *p* = 0.14). We were unable to perform subset analyses for the different oxidative stress components and telomere dynamics due to the low number of studies in this analysis. Nor did we perform any moderator analyses for the same reason. However, the overall effect size included the effects of both antioxidant and oxidant manipulations, and both contributed to the overall effect size.

**Figure 3.**
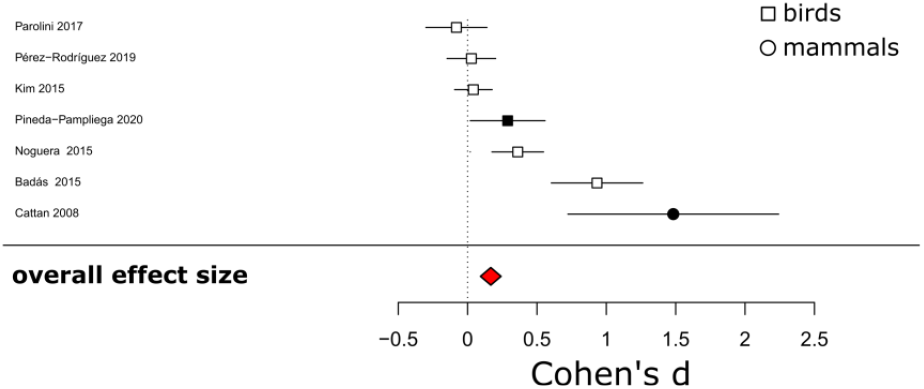
Forest plot of experimental studies manipulating oxidative stress and their Cohen’s d effect sizes on telomere dynamics. Positive effect sizes support the hypothesis that oxidative stress elevation causes telomere attrition (see methods for details on effect size direction). Squares and circles indicate birds and mammals, respectively. The effect sizes in black represent studies measuring telomere length with TRF, whereas the other studies used qPCR. Horizontal bars show the 95% confidence intervals.

## Discussion

We tested the fundamental hypothesis that oxidative stress causes telomere attrition *in vivo*. Our meta-analysis shows that the overall correlation between oxidative stress and telomere dynamics is indistinguishable from zero, questioning this hypothesis. However, we found that the method with which studies measured telomere length significantly affected the meta-analysis (Fig. 2). The subset of studies that measured telomere length with TRF (n=8) showed a small but significant correlation between oxidative stress and telomere dynamics, while the larger subset of qPCR-based studies (n=29) failed to detect a relationship. We attribute this difference to the lower repeatability inherent to the qPCR method (Kärkkäinen et al. 2021) explaining a weaker correlation for the subset of studies that measured telomere dynamics with qPCR. It is therefore likely that the TRF studies more accurately reflect the true biological association between oxidative stress and telomere dynamics *in vivo*. Furthermore, our meta-analysis of experimental studies show a clear effect of treatments alleviating or elevating oxidative stress on telomere dynamics (Fig. 3). We conclude that there is a small but profound effect of oxidative stress on telomere dynamics, but that methodological challenges make it hard to detect this association in living organisms.

Our findings raise the prospect that ageing is underpinned by oxidative stress effects on telomere attrition. However, it remains challenging to functionally link the effect of oxidative stress on telomeres to ageing. Our moderator analyses may shed light on this; if oxidative stress causes both telomere attrition *and* ageing, then we would expect a more profound association between oxidative stress and telomere dynamics in faster ageing organisms. We found a significant negative effect of species longevity (Fig. 2), indicative of a stronger correlation between oxidative stress and telomere dynamics in faster ageing species. Although this finding matches our prediction, it should be noted that the same pattern could emerge when a third unknown factor causes both ageing and oxidative stress. We also found a stronger correlation for the adult life-phase during which ageing becomes apparent (Fig. 2). On the one hand this finding is counterintuitive given that the rate of telomere attrition is generally higher during development (Zeichner et al. 1999; Salomons et al. 2009). On the other hand, negative effects on telomere length could progressively accumulate with age if oxidative stress is moderately consistent through the life course, assuming negligible telomere repair. Following this logic, telomere attrition would be particularly revealing of oxidative stress during the early life phase whereas telomere length would be more revealing late in life. We included the interaction between telomere dynamic and life phase to test this prediction, but this interaction term was not significant (p=0.8). It would be important to further investigate the detailed links between oxidative stress, telomere dynamics, and ageing using experimental and longitudinal approaches to unravel the proximate role of oxidative stress and telomere dynamics in ageing. It would also be important to measure telomerase activity in such study because a causal effect of oxidative stress on telomere attrition could be masked by telomere repair.

Another interesting finding of our moderator analysis is that there were no significant effects of the type of oxidative stress marker or telomere dynamic (Fig. 2). We anticipated a stronger association for longitudinal studies measuring oxidative stress effects on telomere attrition, because such within-individual measurements are less confounded by the effects of among individual heterogeneity (Pol & Wright 2009). However, we found a trend in the opposite direction with a stronger association for studies that measured telomere length (Fig. 2). This is a curious finding for which we have no explanation yet. We found no significant effect of taxonomic group, suggesting that the association between oxidative stress and telomere dynamics could be universal in birds and mammals. It would be interesting to test the effect of oxidative stress on telomere dynamics in ectotherms and aquatic species as these groups are currently lacking.

Even though our moderators substantially reduced the observed study heterogeneity there was significant residual variation indicating that there are other factors involved that we did not capture in our meta-analysis. Interspecific variation in the heritability of telomere length could be important (Chik et al. 2022). Genetic variation appears much smaller for telomere attrition (Bauch et al. 2021) but there can be many other non-genetic factors causing variation in telomere attrition. For example, telomere attrition may be linked to key life-history variables such as growth rate that are themselves not necessarily related to oxidative stress. In the case of growth, this could be mechanistically underpinned by glucocorticoids affecting mitochondrial metabolism and the rate of telomere attrition (i.e. the metabolic telomere attrition hypothesis (Casagrande & Hau 2019)), independently of oxidative stress, as recently observed in great tit nestlings (Casagrande et al. 2020). The oxidative stress and metabolic telomere attrition hypotheses are not mutually exclusive but could obfuscate the predicted relationships for each hypothesis, demanding experimental studies to untangle their independent effects. The residual heterogeneity could also reflect that oxidative stress is difficult to measure. For example, blood parameters are an indirect measurement of the oxidative stress that is occurring within cells and hence a lack of a correlation with telomere attrition could simply mean that the marker does not accurately reflect intracellular oxidative stress. The experimental studies on the other hand show an effect of treatment that does not rely on measurements of oxidative stress. We attempted looking at different types of oxidative stress markers in our meta-analysis to determine if some markers could be more revealing, but we found no evidence for this.

The estimate of our meta-analysis reflects the overall effect size on the organism level from parameters measured in the blood, but there may be large variation among different components of oxidative stress physiology and across differing organs and tissues (Cattan et al. 2008). For example, in humans, the relationship between oxidative stress and telomere attrition appears particularly evident in ovarian tissues (Yang et al. 2021). Other tissues may be better protected depending on their functional importance for immediate survival (Boonekamp et al. 2018). Such prioritization of function could differ between sexes, populations, environments, and species.

Longitudinal studies elevating oxidative stress may shed light on these questions, but it is challenging to perform such experiments in natural populations, which is pivotal for untangling the fitness consequences. Moving forward, it could be possible to exploit altitudinal variation as a natural laboratory setting to address these challenges. Oxidative stress levels depend on altitude with high altitude disrupting antioxidant protection elevating oxidative stress despite the lower partial oxygen pressure (Dosek et al. 2007; Jefferson et al. 2004). Different patterns for oxidative stress across altitudes are found for different species (Dupoué et al. 2017; Stier, Delestrade, et al. 2014; Zhang et al. 2015). Experimental translocations across altitudes could be an interesting way forward to uncover the relationships between the environment, oxidative stress, and the entwined telomere and life history dynamics. In the laboratory, it would be possible to use a more direct approach by manipulating oxygen pressure as previously suggested (Schantz et al. 1999). We hope that such experiments eventually further our understanding of the proximate causes of ageing.

## Materials and Methods

### Data collection

We performed a literature search on the Web of Science using the search string “Telom* AND Oxidat* OR Telom* AND Antioxi*” ending on 28/01/2022. A total of 5,526 of papers from both human and animal studies were screened (Fig. 4). Additional papers were found by screening the cited papers in Reichert and Stier (2017) and Boonekamp *et al*. (2017) and by checking the relevant citations from these papers. We exclusively used studies on healthy non-experimental subjects. Where necessary we approached the authors for additional information on the healthy control population, for example when the reported findings were on the combined healthy control and diseased or experimental subjects. We did not include studies using 8-OHdG, measured in urine, as a marker of oxidative damage (n=6). 8-OHdG are oxidized guanine base pairs generally interpreted to reflect DNA damage (Wu et al. 2004). However, 8-OHdG compounds are only cleaved when DNA repair occurs, and our interpretation is therefore that 8-OHdG secretion is a function of both damage and repair. Hence, 8-OHdG is an interesting but fundamentally different biomarker than the other markers of oxidative stress that were included in our meta-analysis.

**Figure 4:**
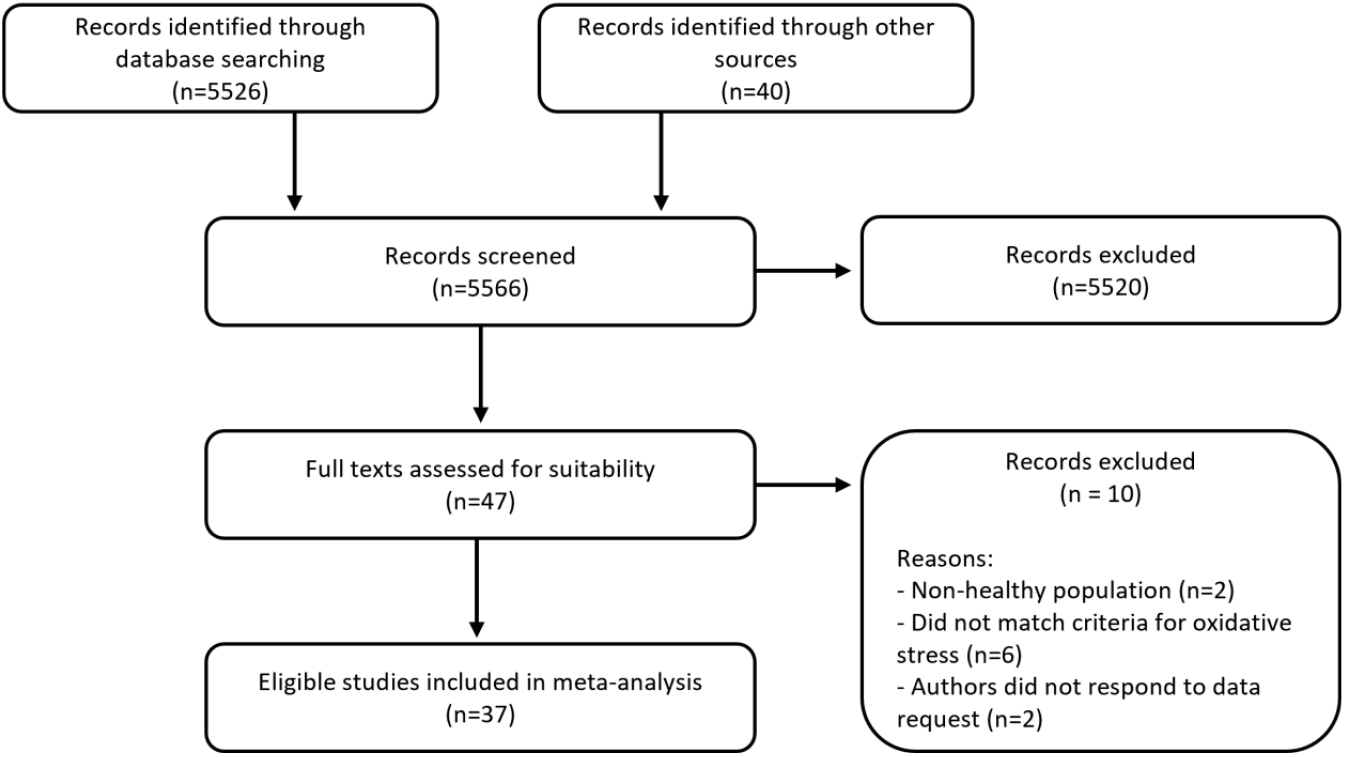
Preferred Reporting Items for Systematic Reviews and Meta-Analyses (PRISMA) flow diagram of studies checked for eligibility in our meta-analysis. The initial search was done through key word searching in ISI-Web of knowledge and further potentially suitable records were found through scanning the references and citations of two key papers in the field (see methods for details).

Overall, we determined 107 Pearson correlations from 37 papers on the associations between markers of oxidative stress and telomere dynamics. We used the reported Pearson correlations when available, or the reported *p*-values and sample sizes to calculate a t-statistic which we then converted into a Pearson correlation effect size following equation 11 in (Nakagawa & Cuthill 2007). When correlations were reported for associations between oxidative stress and telomere attrition in addition to telomere length, we only used the estimates for telomere attrition as we assumed these would be more accurate, being less affected by individual variation in baseline telomere length.

Our search yielded 7 additional studies that experimentally manipulated oxidative stress investigating a causal relationship between oxidative stress and telomere attrition. This is a fundamentally different type of study estimating the mean telomere difference between control and treatment groups, which is best expressed in the form of a standardized mean difference, or Cohen’s *d* effect size. We conducted an independent meta-analysis for this subset of studies by determining the Cohen’s *d* effect sizes using the reported mean telomere differences and standard deviations between treatment and control groups (following equation 1 in Nakagawa & Cuthill 2007). When the mean and standard deviations were unavailable, we calculated a Pearson correlation (using reported *p* and *n* statistics) which we then transformed into a Cohen’s *d* estimate using equation 26 in (Nakagawa & Cuthill 2007). Table 1 provides an overview of the studies that were included in our meta-analyses. Data table S1 shows the full overview of the effect sizes that were extracted from these studies.

### Meta-analysis and directionality of effect sizes

We used the metafor package (v 3.4.0; (Viechtbauer 2010)) in R (version 4.0.3) to conduct a multilevel mixed effects model with study ID as random effect (Viechtbauer 2010), accounting for the non-independence of multiple effect sizes from the same study population. Sampling variances were determined with the Hunter and Schmidt method for correlational studies (Schmidt & Hunter 2015) and we used the inversed variances for Cohen’s *d* effect sizes for experimental studies (29). We used the Kendall’s tau statistic to test for publication bias and Q-tests for evaluating effect size heterogeneity. We attempted to investigate the potential causes of heterogeneity by including methodologically and biologically relevant moderators. Two commonly used telomere length measurement methods are Terminal Restriction Fragment (TRF) analysis and quantitative PCR (qPCR). These methods differ in their technical and within-individual repeatability, with TRF being more accurate than qPCR (Verhulst et al. 2015; Kärkkäinen et al. 2021). We have previously shown that there are systematic differences in the association between telomere length and mortality amongst TRF- and qPCR-based studies (Wilbourn et al. 2018). We therefore included telomere length measurement method (qPCR vs. TRF) as a moderator variable in our meta-analysis. We included taxonomic class (Aves vs. Mammalia), life phase (pre- vs. post-reproductive maturation), and the logarithm of maximum longevity (using the species information available on “AnAge”:https://genomics.senescence.info/species/index.html)as additional moderator variables. Moderators’ significance was assessed using z-tests. We report all effect sizes with their 95% confidence intervals (CI).

To estimate the overall relationship between oxidative stress markers and telomere dynamics all effect sizes were pooled into a single meta-analysis; Both the expected positive relationships (i.e. oxidative damage and telomere attrition; antioxidant protection and telomere length) and negative ones (i.e. oxidative damage and telomere length; antioxidant protection and telomere attrition) supporting the underlying hypothesis that oxidative stress causes telomere attrition were pooled into a single meta-analysis. To facilitate a unidirectional interpretation, we flipped the sign of the expected negative relationships (oxidative damage and telomere length; antioxidant protection and telomere attrition) *meaning that all positive effect sizes in our meta-analysis support the hypothesis that oxidative stress increases telomere attrition*. This approach also enabled us to formally test if the association between oxidative stress and telomere dynamics depended on the type of oxidative stress marker (damage vs. protection), or telomere dynamic (telomere length vs. telomere attrition) by including these variables as additional moderators in the meta-analysis.

## Acknowledgments

We thank Agnes Saulnier, Antoine Stier, Helena Schmidt and Jing Shen for providing additional information and/or data, enabling including their studies in our meta-analysis.

## Author Contributions

This study was conceived by JJB. EA performed the literature search, extracted the relevant statistics calculated effect sizes, and performed the initial meta-analysis. JJB curated the final dataset, performed the final meta-analyses, and wrote the first draft of the manuscript. Both authors contributed to subsequent revisions.

## Data availability statement

Study meta-information, effect sizes, and moderator variables that were used in this meta-analysis will be made available on Dryad upon acceptance for publication.

